## Supplementary information for "Cell autonomous role of leucine-rich repeat kinase in protection of dopaminergic neuron survival"

### Figure 1-figure supplement 1. Generation of the *LRRK1* targeting vector

**1)** The *left middle* arm was amplified by PCR from mouse BAC DNA (Clone #: RP23-213J23, BACPAC Resources Center) using primers P3 and P4. The resulting PCR product (2,079 bp) containing part of *LRRK1* intron 26, exon 27, and part of intron 27 was subcloned into the pGEM-T Vector (Promega, Cat#; A1360) by TA ligation to generate pLRRK1#1 (pLM1).

P3: 5'CCAGTCACTTCTCCACCTCAGGGAAAATGG (30 bp)

P4: 5'CCTTGT*GGTACC*CGGACCTTCTATCACCTTTATCC (35 bp): *Kpn*I (*GGTACC*) is an endogenous restriction site.

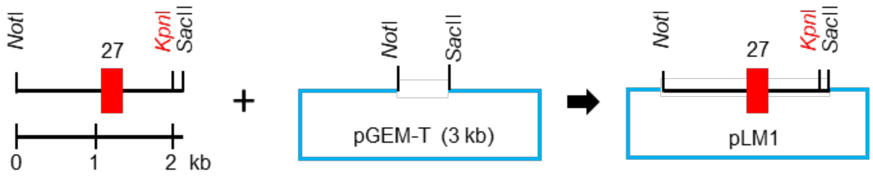


**2)** The *right middle* arm was amplified from mouse BAC DNA (Clone #: RP23-213J23) using primers P5 and P6. The resulting PCR product (3,403 bp) containing part of *LRRK1* intron 27, exon 28, intron 28, exon 29, and part of intron 29 was subcloned into the pGEM-T Vector by TA ligation to generate pLRRK1#2 (pLM2).

P5: 5'GGTCCG*GGTACC*ACAAGGTGCTGGTTAAGTGCC (33 bp): *Kpn*I (*GGTACC*) is an endogenous restriction site.

P6: 5'AGCAGACCTCTTGCCTTCTACTACTGACTG (30 bp)

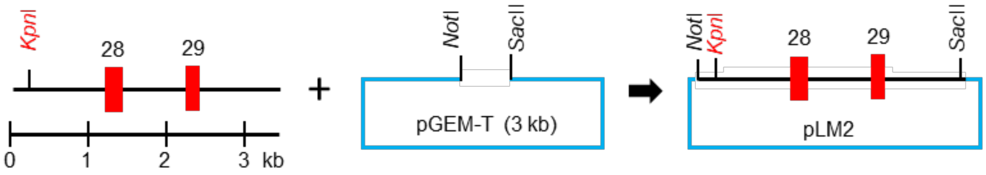


**3)** pLRRK1#1 (pLM1) was digested with *Not*I and *Kpn*I, then subcloned into the *Not*I and *Kpn*I sites of pLRRK1#2 (pLM2) to generate the pLRRK1#3 (pLM3) plasmid which contains the *middle* homologous region.

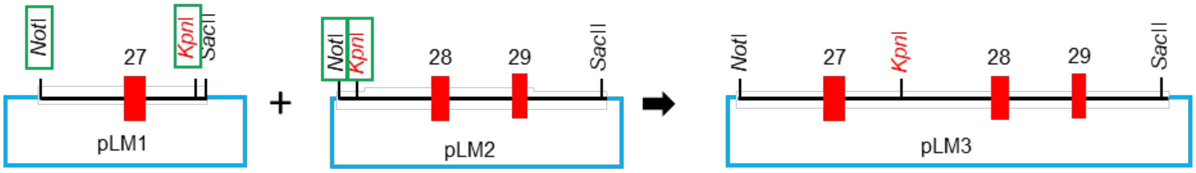


**4)** The middle homologous region was released from pLRRK1#3 (pLM3) using *Not*I and *Sac*II followed by Klenow to blunt the sticky ends, which was then subcloned into the *Sma*I site of pGKneoF2L2DTA (addgene: #13445) to generate pLRRK1#4 (pLM4).

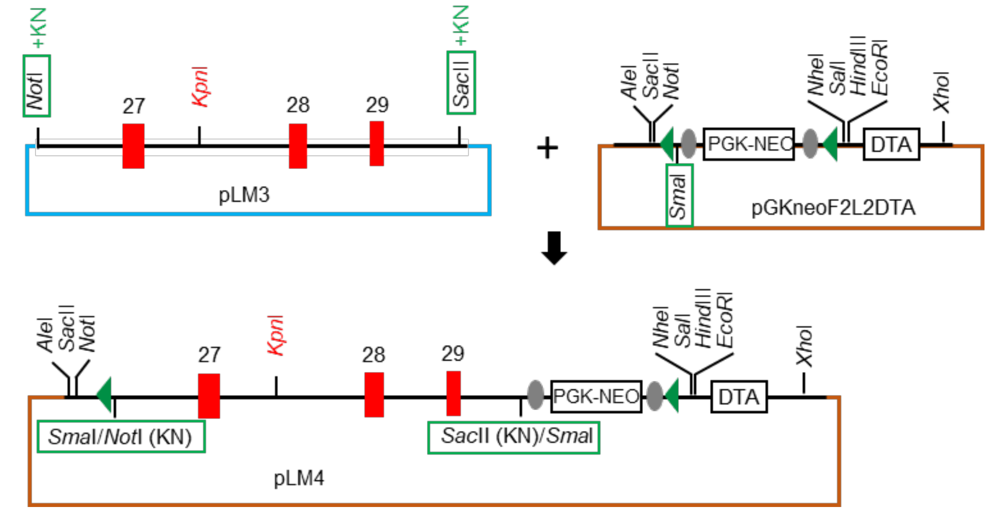


Green arrowhead: *loxP* site, gray circle: *FRT* site

**5)** The left arm was amplified from mouse BAC DNA (Clone: RP23-213J23) using primers P1 and P2. The resulting PCR product (2,016 bp) containing part of *LRRK1* intron 25, exon 26, and part of intron 26 was digested with *Sac*II and *Not*I, and was then subcloned into the *Sac*II and *Not*I sites of pGKneoF2L2DTA (addgene: #13445) to generate pLRRK1#5 (pLM5).

P1: 5'gacat*CCGCGG*CACCATGTGAGTGGCAGCTGTGGTGAGAAC (41 bp). *Sac*II (*CCGCGG*) is an exogenous site.

P2: 5'gacat*GCGGCCGC<u>AAGCTT</u>*TTTAATAGCCGTTCTTTCTTAGAGAAGGCAG (50 bp). *Not*I (*GCGGCCGC*) and *Hind*III (*<u>AAGCTT</u>*) are exogenous sites.

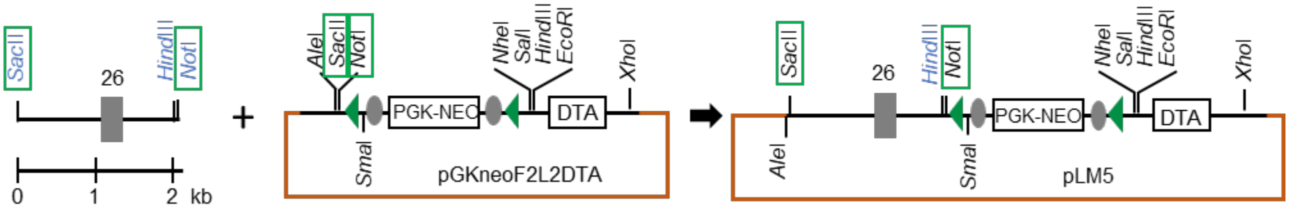


**6)** The right arm was amplified from mouse BAC DNA (Clone: RP23-213J23) using primers P7 and P8. The resulting PCR product (3,131 bp) containing part of *LRRK1* intron 29, exon 30, and part of intron 30 was digested with *Sal*I and *Hind*III, and was then subcloned into the *Sal*I and *Hind*III sites of pGKneoF2L2DTA (addgene: #13445) to generate pLRRK1#6 (pLM6).

P7: 5'gacat*GTCGAC*GGATCCGTAGGGAAGACCCACTAGGAGGAAGAAAG (46 bp). *Sal*I (*GTCGAC*) is an exogenous site.

P8: 5'gacat*AAGCTT*TGGTACCTTTCTAAAGGCAGCATTTTGCTTGC (43 bp). *Hind*III (*AAGCTT*) is an exogenous site.

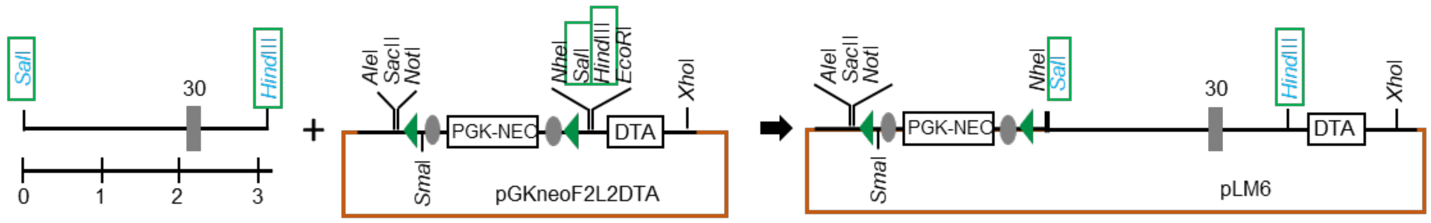


**7)** pLRRK1#5 (pLM5) was digested with *Sac*II and *Not*I, and was then subcloned into the *Sac*II and *Not*I sites of pLRRK1#6 (pLM6) to generate pLRRK1#7 (pLM7).

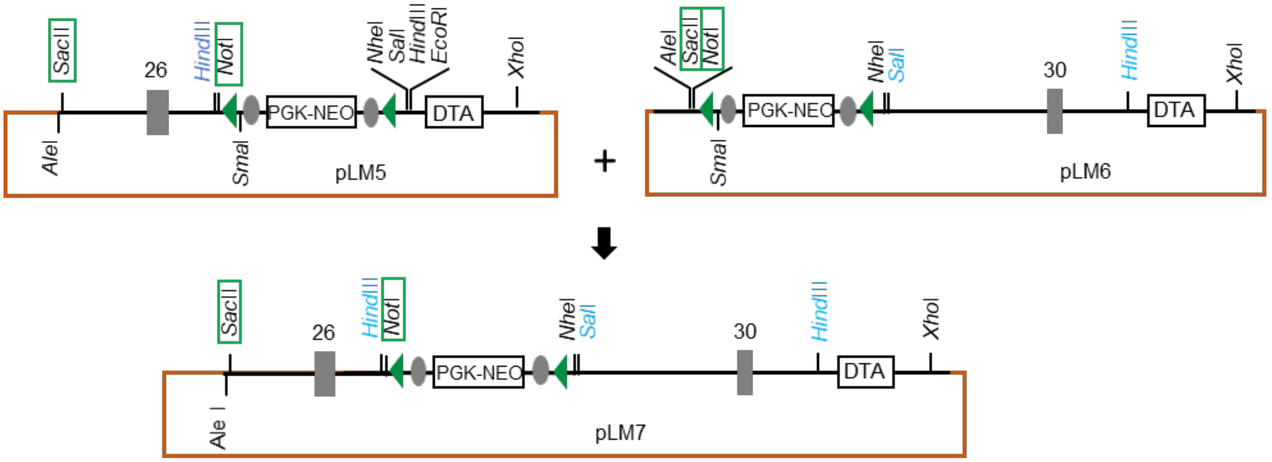


**8)** The middle homologous region was released from pLRRK1#4 by *Not*I and *Sal*I and was then subcloned into the *Not*I and *Sal*I sites of pLRRK1#7 (pLM7) to generate the final targeting vector pLRRK1#8 (pLM8), which was linearized with *Xho*I before electroporation into ES cells.

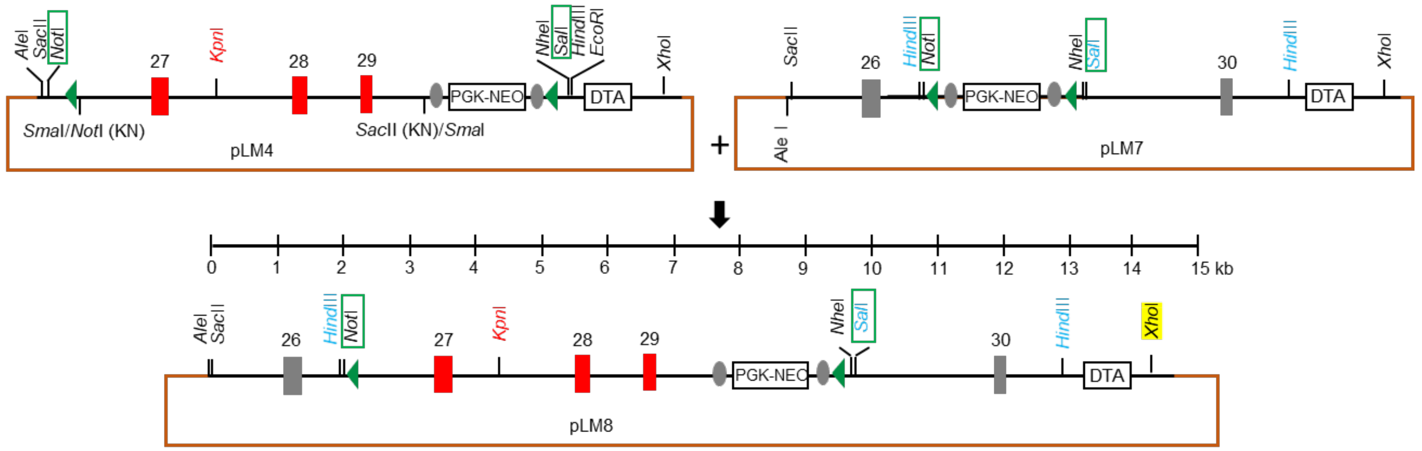


Restriction sites in black: from vectors

Restriction sites in red: *LRRK1* endogenous sites

Restriction sites in blue: introduced by the primers

**Figure 1-figure supplement 2. Genomic DNA sequence of the floxed *LRRK1* allele**

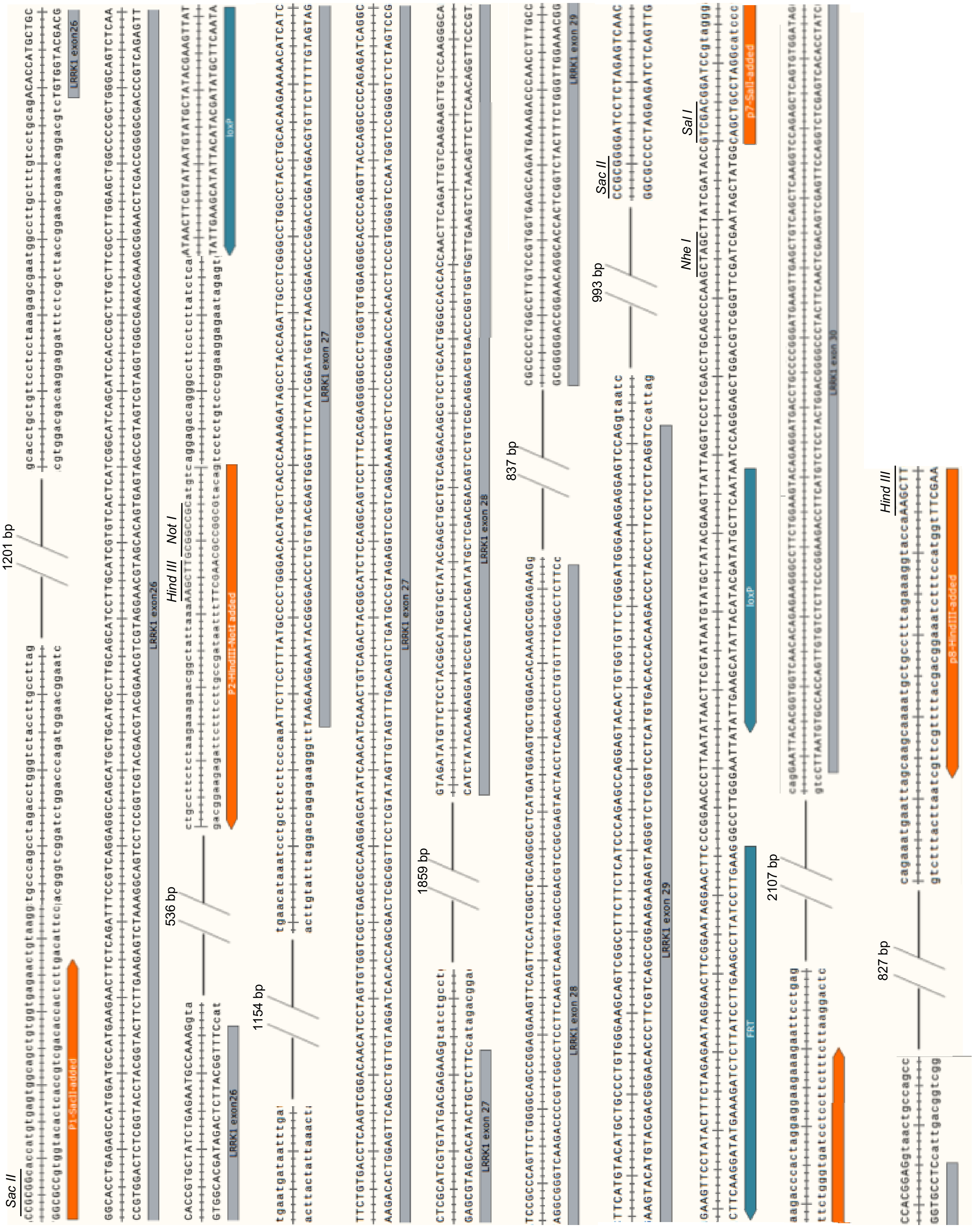

### Figure 1-figure supplement 3. Southern strategy for the targeted and floxed *LRRK1* alleles

**5' external probe:** 377 bp (*LRRK1* intron 25)

P15: 5'CAGAATACCCCATGCTGGGGAATTGC (26 bp)

P16: 5'CCGTTTCTAGGATTCTAATTTTTC (24 bp)

***Hind III digestion*** ***WT: 17.0 kb***
***Targeted: 4.8 kb***
***Floxed: 4.8 kb***

**3' external probe:** 305 bp (*LRRK1* exon 32)

P17: 5'AACTCCTTCCTGGTGCTGGCAGGCCTGGCTG (31 bp)

P18: 5'ACAAGTGACGTGACCATGGACGGAGCTGCG (30 bp)

***Hind III digestion*** ***WT: 17.0 kb***
***Targeted: 14.1 kb***
***Floxed: 12.2 kb***

***Neo*** **probe (363 bp)**

P23: 5'ATTCGGCTATGACTGGGCAC (20 bp)

P24: 5'GACCACCAAGCGAAACATCG (21 bp)

***Hind III digestion*** ***WT and Floxed: no band***
***Targeted: 14.1 kb***

### Figure 1-figure supplement 4. Generation of the *LRRK2* targeting vector

**1)** The left homologous region was amplified from mouse BAC DNA (Clone# RP23-526-A2, BACPAC Resources Center) using primers P9 and P10. The resulting PCR product (2,579 bp) was digested with *EcoR*I, which is an endogenous restriction site upstream of the *LRRK2* promoter, and was then subcloned into the *EcoR*I and *Sma*I sites of pBSK (+) to generate pLRRK2#1 (2,526 bp + 2,961 bp = 5,487 bp).

P9: 5' GAACACACAAGGCTATGGCTATTGTC (26 bp)

P10: 5' GTAGGACTATCATCCACCTGTAGGACTCC (29 bp)

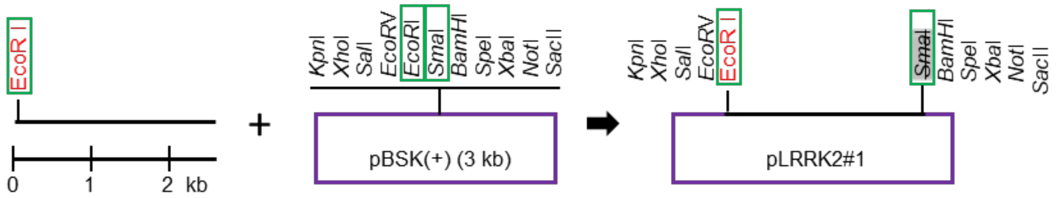


**2)** Oligoes (P39 and P40) were annealed together to generate a double stranded polylinker (*BamH*I-*loxP*-*Nhe*I-*Spe*I), which was then subcloned into the *BamH*I and *Spe*I sites of pLRRK2#1 to generate pLRRK2#2 (5,487 bp + 50 bp = 5,537 bp).

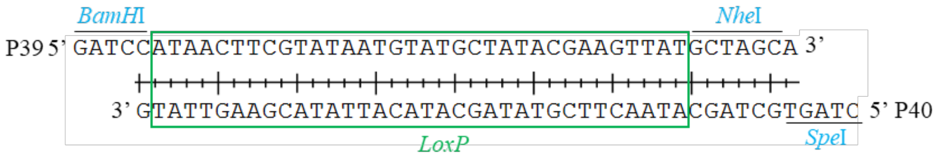


P39: 5' GATCCATAACTTCGTATAATGTATGCTATACGAAGTTATGCTAGCA (46 bp).
*BamH*I (G'GATCC), *loxP* top strand (ATAACTTCGTATAATGTATGCTATACGAAGTTAT), *Nhe*I (G'CTAGC).

P40: 5' CTAGTGCTAGCATAACTTCGTATAGCATACATTATACGAAGTTATG (46 bp).
*Spe*I (A'CTAGT), *Nhe*I (GCTAGC), *loxP* bottom strand (ATAACTTCGTATAGCATACATTATACGAAGTTAT).

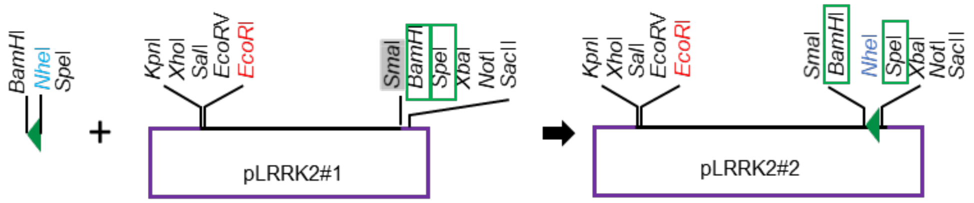


Green arrowhead: *loxP* site

**3)** The middle arm was amplified from mouse BAC DNA (Clone# RP23-526-A2) using primers P11 (upstream of the *LRRK2* promoter) and P12 (intron 2). The resulting PCR product (2,990 bp) was digested with *Xba*I and *Not*I and was then subcloned into the *Xba*I and *Not*I of pGEM-T Vector (Promega, Cat#: A1360) to generate pLRRK2#3 (2,990 bp + 3,000 bp = 5,990 bp). *Xba*I, *Not*I, and *Nhe*I sites are exogenous sequences introduced into PCR primers.

*Xba*I

PCR: P11=5' GTCCTACAGGTGGATTCTAGACCTACAAGG 3' (30 bp)

P12=5' GACGCGGCCGCGTTTAGTAGCTAGCATGACTATGAAGGAG 3' (40 bp)

*Not*I *Nhe*I

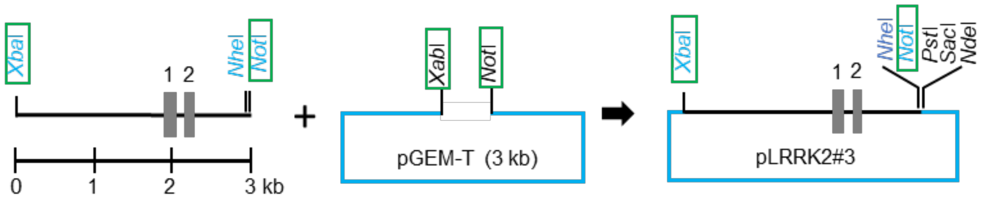


**4)** pLRRK2#3 was digested with *Xba*I and *Not*I, and the insert was then subcloned into the *Xba*I and *Not*I sites of pLRRK2#2, which contains the left homologous arm and the *loxP* sequence, to generate pLRRK2#4 (2,958 bp + 5,490 bp = 8,448 bp).

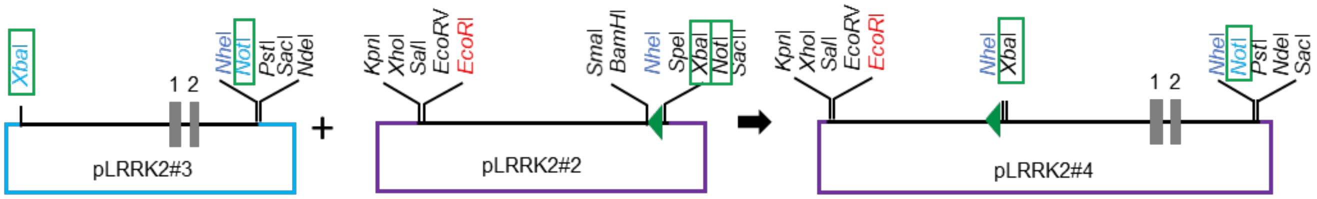


**5)** The right arm was amplified from mouse BAC DNA (Clone# RP23-526-A2) using primers P13 and P14. The resulting 3,503 bp PCR fragment containing *LRRK2* intron 1, exon 2, and intron 2, was subcloned into the *EcoR*V site of pBSK(+) to generate pLRRK2#5 (3,503 bp + 2,961 bp = 6,464 bp). **Check direction!!!**

PCR: P13=5'GCACTTGAGTCTTAATCTTGGGCAC 3' (25 bp)

P14=5'CATTCGAGCAGCTAAGCCTGTAATC 3' (25 bp)

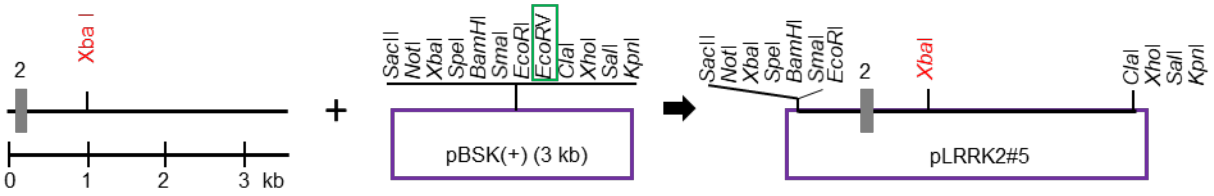


**6)** pLRRK2#5 was digested with *BamH*I and then treated with Klenow to blunt the overhang followed by *Cla*I digestion. The digested fragment was subcloned into the *EcoR*V and *Cla*I sites of the pSoriano vector (*PGKneolox2DTA*, addgene 13443) to generate pLRRK2#6 (3,547 bp + 6,307 bp = 9,854 bp).

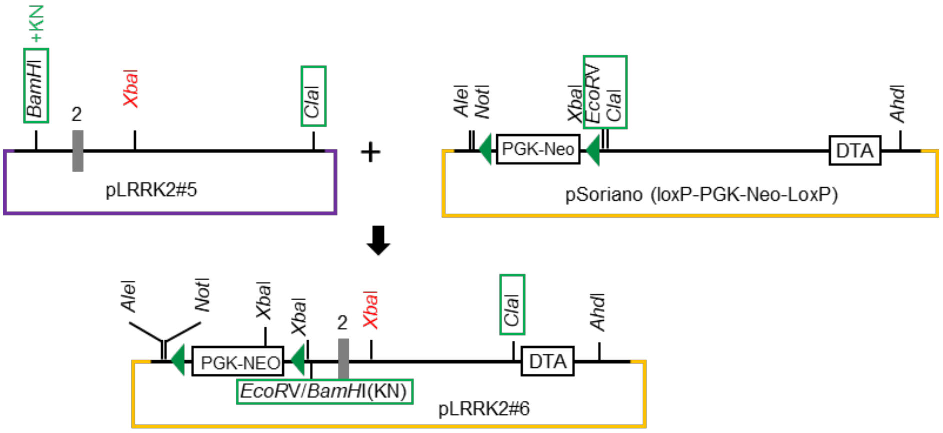

**7)** The LFNT-tk/pBS plasmid (a gift from S. Tonegawa) was digested with *Sac*II (followed by Klenow) and *Not*I (followed by *Ssp*I) to release the "*loxP-FRT-PGK-neo-loxP-FRT*" fragment (2,928 bp), which was then subcloned into the *Xba*I (followed by Klenow) and *Not*I sites of pLRRK2#6 to generate pLRRK2#7 (2,928 bp + 7,051 bp = 9,979 bp).

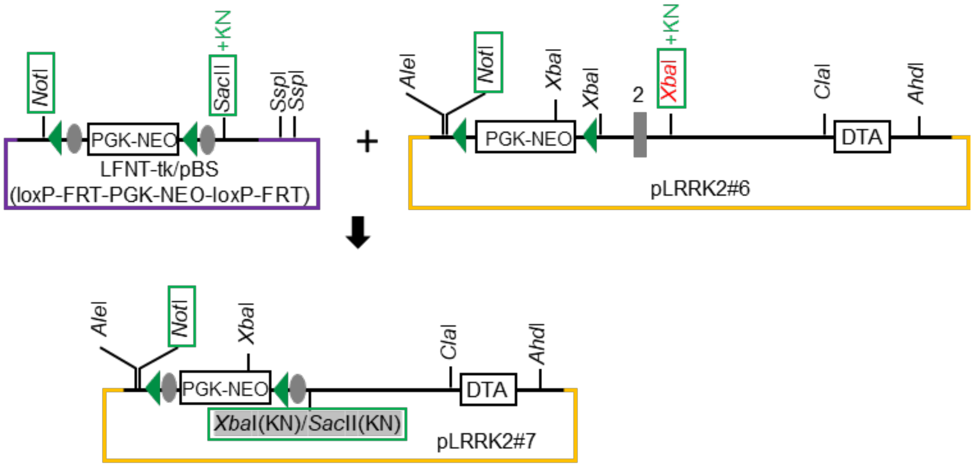


Green arrowhead: *loxP* site, gray circle: *FRT* site

**8)** pLRRK2#4 was digested with *Eco*RV and *Not*I to release the fragment containing the left arm-*loxP*-middle arm, which was then subcloned into the *Ale*I and *Not*I sites of pLRRK2#7 to obtain pLRRK2#8, the *LRRK2* targeting vector (15,485 bp). The linearized targeting vector (with *Ahd*I) was electroporated into ES cells.

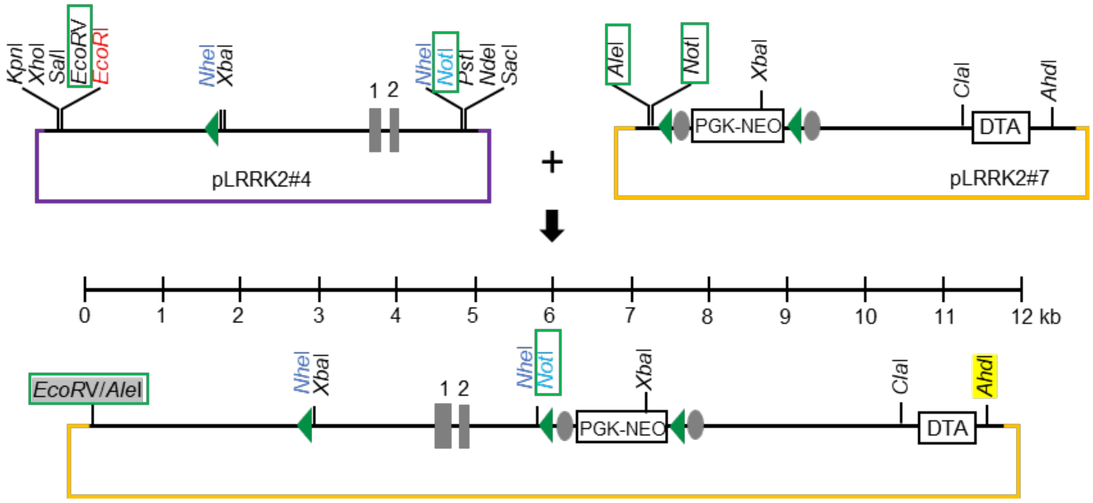


Restriction sites in black: from vectors

Restriction sites in red: *LRRK2* endogenous sites

Restriction sites in blue: introduced by the primers

**Figure 1-figure supplement 5. Genomic DNA sequence of the floxed *LRRK2* allele**

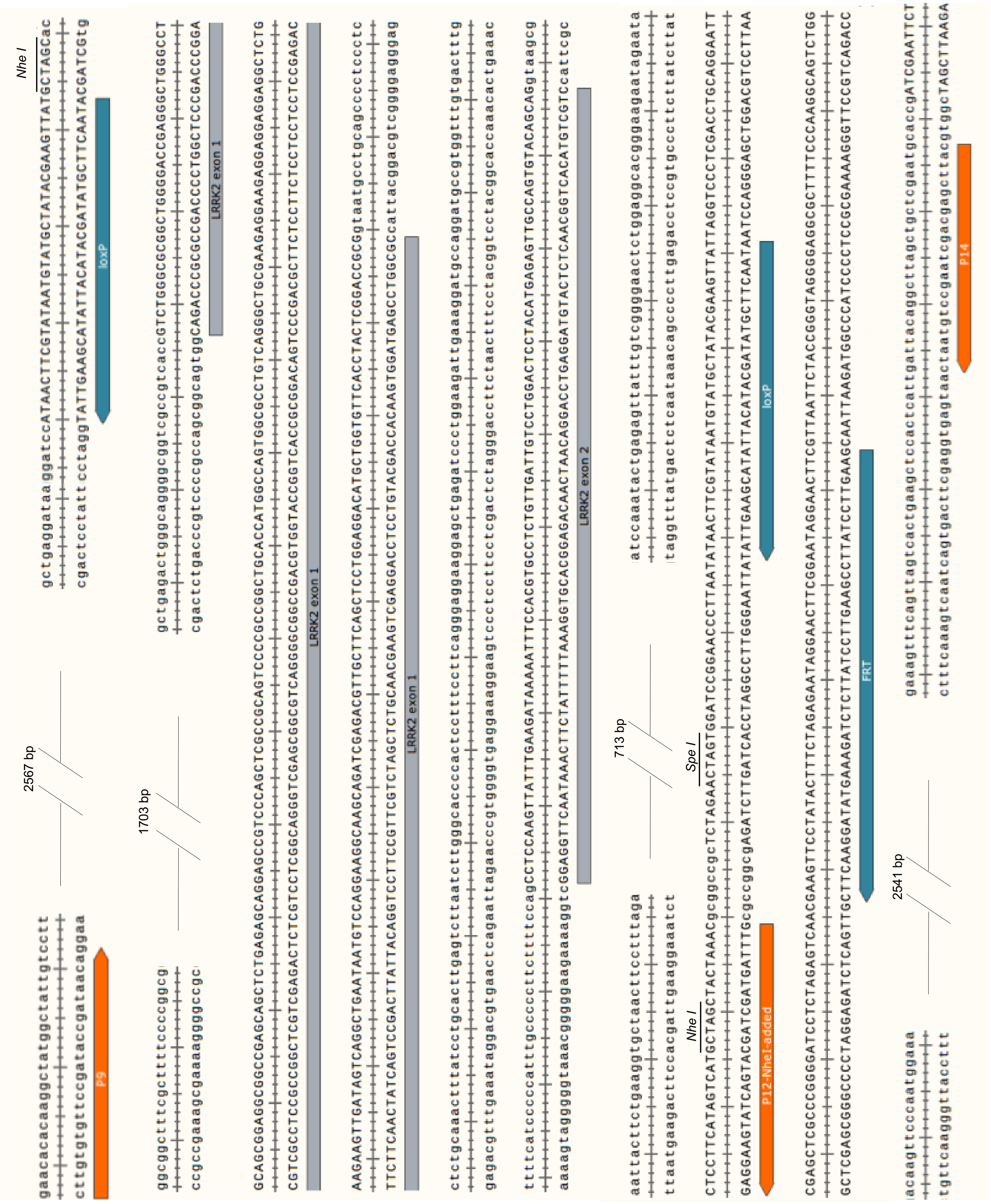

### Figure 1-figure supplement 6. Southern analysis of the targeted and floxed *LRRK2* alleles

**5' external probe (753 bp)**

P19= 5'CATGGGAGAGAGGGTTTCTCACTTACT (27 bp)

P20= 5'CTTGGACAGCATTGTCAGCCTAGAC (25 bp)

| | |
|---|---|
| ***Nhe I digestion*** | ***WT: 11.5 kb*** <br> ***Targeted and floxed: 3.6 kb*** |

**3' external probe (622 bp)**

P21=5' GCCAAGCAGTTATTGATGCTGTAGC (31 bp)

P22= 5' CGAGCTGTAAGATGAGCTGGGTACT (30 bp)

| | |
|---|---|
| ***Sph I digestion*** | ***WT: 12.3 kb*** <br> ***Targeted: 4.5 kb*** |
| ***Nhe I digestion*** | ***WT: 11.5 kb*** <br> ***Floxed: 5.2 kb*** |

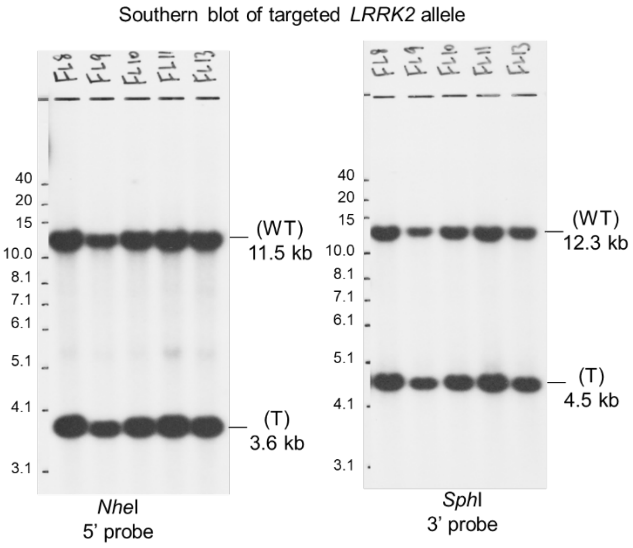

### Figure 1-figure supplement 7. Northern strategy for *LRRK1* mRNA

***LRRK1* exons 2-3 probe (377+4=383 bp)**

P31: 5' CAGGATGAGCGTGTGTCTGCAG (CAG exon 1/rest exon2)

P32: 5′ CCTTCTCCTGTGAGGATTCGCTCT (C exon 3/rest exon 4)

***WT and floxed alleles: ~7.4 kb***

***Deleted allele: ~6.8 kb****

* Deletion of *LRRK1* exons 27-29 (625 bp; exon 27: 278 bp, exon 28: 195 bp, exon 29 152 bp) results in frameshift of downstream exons, likely resulting in degradation of the transcript.

***LRRK1* exons 27-29 probe (550 bp)**

P33: 5' CTGGCCTACCTGCACAAGAA (exon 27)

P34: 5' CCTTCCCATCCCAGAACACC (exon 29)

***WT and floxed alleles: 7.4 kb***

***Deleted allele: no band***

***GAPDH* probe (452 bp)**

P37: 5' ACCACAGTCCATGCCATCAC (exon 5)

P38: 5′ TCCACCACCCTGTTGCTGTA (exon 7)

**Figure 1-figure supplement 8. Northern analysis of *LRRK1* mRNA in mice carrying *LRRK1* germline deleted alleles**

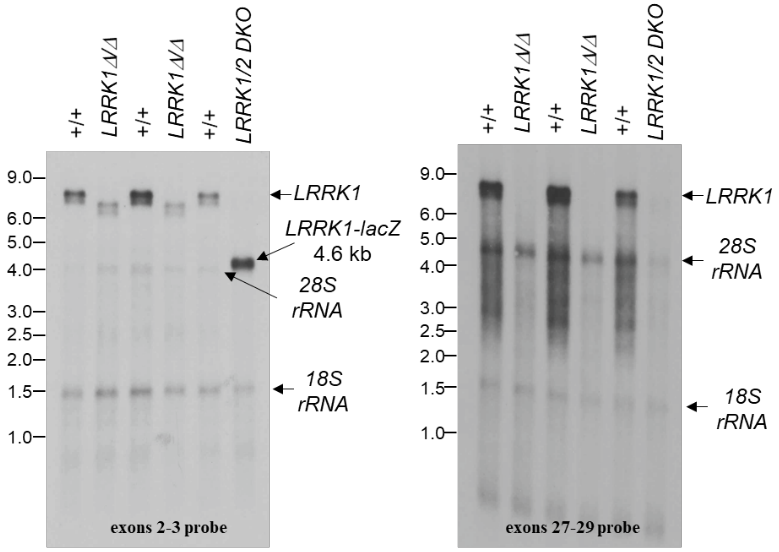


*LRRK1* Northern blots (polyA+ RNA from the lung) using exons 2-3 or exons 27-29 probe: The exons 2-3 probe detected the full-length *LRRK1* transcripts (~7.4 kb) in wild-type (+/+) mice and reduced size transcripts in *LRRK1* Δ/Δ mice, likely due to the absence of exons 27-29 (625 bp), which was confirmed by sequencing of RT-PCR products (see Figure1-figure supplement 9). The truncated *LRRK1* mRNA lacking exons 27-29 is present in lower levels, likely due to mRNA degradation by nonsense-mediated decay, as the absence of exons 27-29 (625 bp) results in a frameshift of downstream codons.

The exons 2-3 probe also recognizes the fusion transcript *LRRK1-lacZ* (~4.6 kb) in *LRRK1/2* DKO mice, which contains *LRRK1* exons 1-3 and exon 2 of engrailed, *IRES* and *lacZ* cDNA (Figure 1, Figure S1 in Giaime et al., *Neuron*, 2017). The exons 27-29 probe also recognizes the full-length *LRRK1* transcripts (~7.4 kb) in wild-type (+/+) mice but no truncated *LRRK1* transcripts, due to the germline deletion of the genomic region from intron 26- intron 29, lacking exons 27-29. The exons 27-29 probe also did not detect the *LRRK1-lacZ* fusion transcript, which only contains exons 1-3, in *LRRK1/2* DKO mice. While polyA enrichment reduced levels of 18S and 28S rRNAs, they are still present in the lanes/samples.

### Figure 1-figure supplement 9. RT-PCR analysis of the deleted *LRRK1* allele

RT-PCR analysis using a *LRRK1* exon 32-specific primer P47 (5'GGCTCAGGTCATGCTCAGTT) for RT and PCR primer sets in *LRRK1* exons 4-8, 11-17, 20-25, and 25-31 indicates normal splicing in homozygous floxed *LRRK1* (F/F) mice and properly deleted transcripts in *LRRK1* Δ/Δ mice. The identity of all PCR products was confirmed by sequencing.

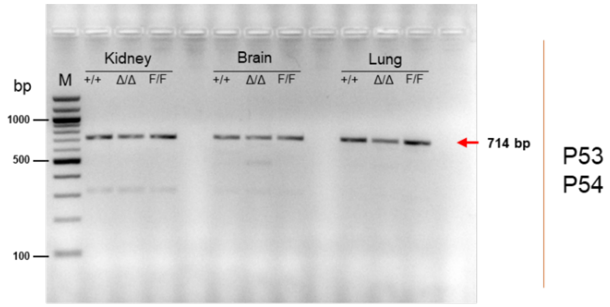


P53: 5' TTTTGGACACGCCGAAGTAGT (**exon 4**)

P54: 5' AGCCGCTCCAGGTAGTTTTT (**exon 8**)

714 bp: Ex4(47)-Ex5(180)-Ex6(149)-Ex7(227)-Ex8(111)

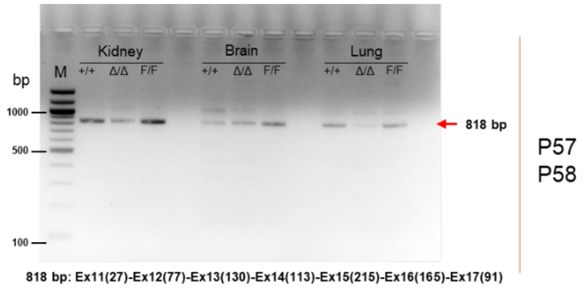


P57: 5' GGACCTCTCCAGAAACCAGC (**exon 11**)

P58: 5' GCAGGGTTGCTATCCTCTCC (**exon 17**)

818 bp: Ex11(27)-Ex12(77)-Ex13(130)-Ex14(113)-Ex15(215)-Ex16(165)-Ex17(91)

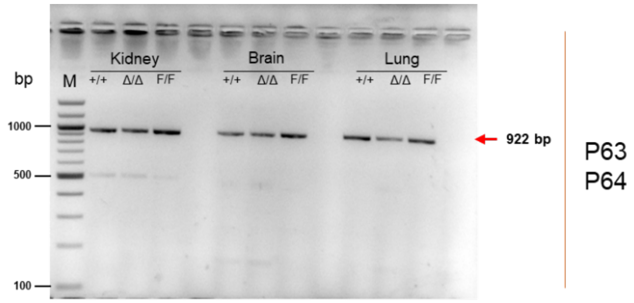


P63: 5' GCGGTCAGTGGCAAAGAATG (**exon 20**)

P64: 5' AATGCTGTTCTCACCCTCCG (**exon 25**)

922 bp: Ex20(144)-Ex21(172)-Ex22(164)-Ex23(140)-Ex24(250)-Ex25(52)

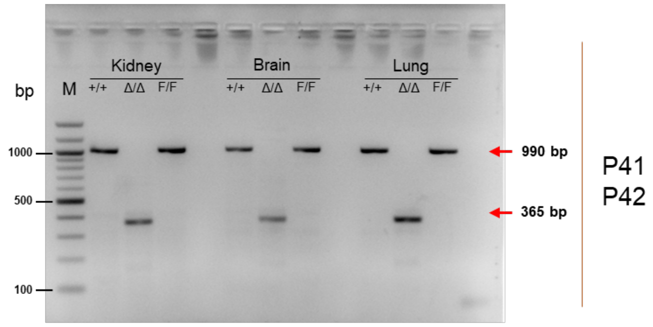


P41: 5' GAATTCTGCTAATGCCCCAGC (**exon 25**)

P42: 5' AGGCTGTAGATATAGATTTTCTGGT (**exon 31**)

The 990 bp product containing exons 25-31 represents the WT allele and confirms the floxed allele. The 365 bp product containing exons 25, 26, 30, and 31 confirms the deletion of exons 27-29.

990 bp: wild-type exon 25 to 31
Ex25(23)-Ex26(198)-Ex27(278)-Ex28(195)-Ex29(152)-Ex30(118)-Ex31(26)
365 bp: deletion of exon 27 to 29
Ex25(23)-Ex26(198)-Ex30(118)-Ex31(26)

**Figure 1-figure supplement 10. Northern analysis of *LRRK2* mRNA in mice carrying floxed or germline deleted *LRRK2* alleles**

***LRRK2* exons 1-5 probe (437 bp)**

P35= 5’ AGGAAGGCAAGCAGATCGAG (exon 1, 20 bp)

P20= 5’ GGCTGAATATCTGTGCATGGC (exon 5, 22 bp)

***WT and floxed alleles: 8.3 kb***
***Deletion: no band***

***GAPDH* probe (452 bp)**

P37= 5’ ACCACAGTCCATGCCATCAC (exon 5, 20 bp)

P38= 5′ TCCACCACCCTGTTGCTGTA (exon 7, 20 bp)

*LRRK2* Northern blot (total RNA from the brain) using exons 1-5 probe: The probe detected the full-length *LRRK2* transcript (~8.3 kb) in wild-type (+/+) and *fLRRK2/fLRRK2* mice, and no transcript in *LRRK2* Δ/Δ mice, derived from floxed *LRRK2* alleles, and previously generated *LRRK1/2* DKO mice (Giaime et al., *Neuron*, 2017).

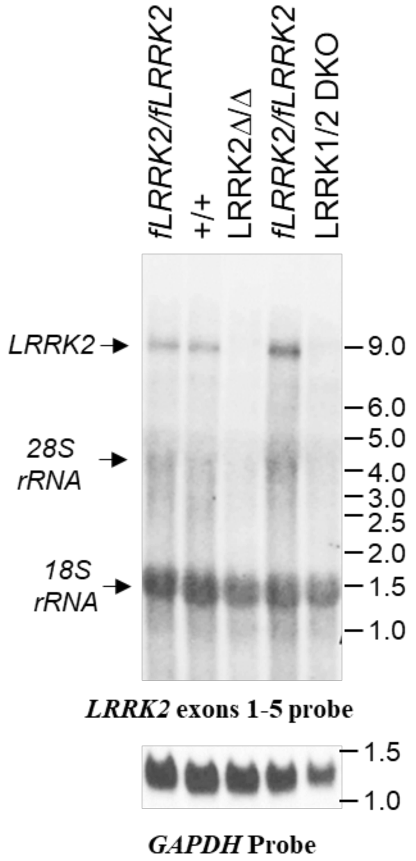

**Figure 1-figure supplement 11. RT-PCR analysis of the deleted *LRRK2* allele**

RT-PCR analysis using an *LRRK2* exon 51-specific primer P93 (5'TCGTGTGGAAGATTGAGGTCC) for RT, and primers in *LRRK2* exons 1 and 5, P35 and P20, for PCR, which shows normal splicing in *fLRRK2/fLRRK2* brains and the absence of PCR products in *LRRK2* Δ/Δ brains. The identity of the PCR product in +/+ and *fLRRK2/fLRRK2* brains was confirmed by sequencing: The 437 bp product contains *LRRK2* exons 1-5.

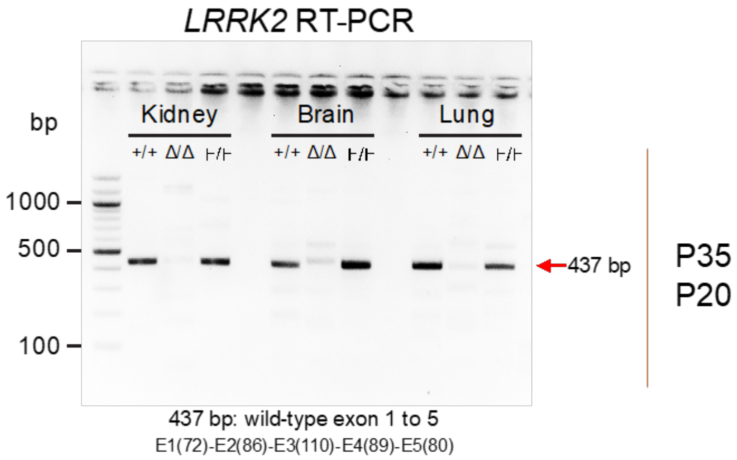


P35= 5' AGGAAGGCAAGCAGATCGAG (exon 1)

P20= 5' GGCTGAATATCTGTGCATGGC (exon 5)

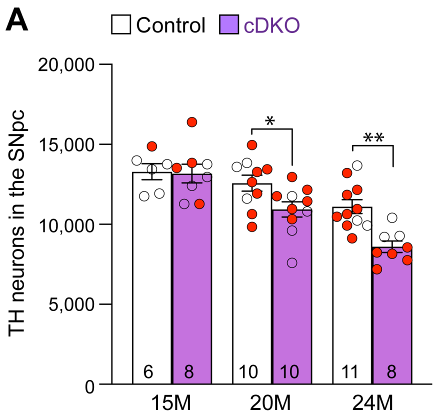


**Figure 3-figure supplement 1. Independent validation of age-dependent reduction of DA neurons in the SNpc of *LRRK* cDKO mice**

**(A)** Quantification of TH+ DA neurons in the SNpc by an independent investigator using stereological methods sampling 25% areas of the SNpc. There are similar numbers of DA neurons in *LRRK* cDKO mice (13,180 ± 585) and littermate controls (13,293 ± 500, $p > 0.9999$) at 15 months of age. At 20 months of age, the number of DA neurons in the SNpc pf *LRRK* cDKO mice (10,936 ± 477) is significantly reduced, compared to control mice (12,576 ± 497, $F_{1,47}=12.40$, $p = 0.0003$; $p = 0.0423$, two-way ANOVA with Bonferroni's post hoc multiple comparisons). By 24 months of age, the reduction of DA neurons in the SNpc of *LRRK* cDKO mice (8,600 ± 355) is more severe, compared to controls (11,105 ± 435, $p = 0.0015$).

The number in the column indicates the number of mice used in the study. Red-filled and open circles represent data obtained from individual male and female mice, respectively. All data are expressed as mean ± SEM. *$p<0.05$, **$p < 0.01$.
