## Supplementary figures and images for "Cell autonomous role of leucine-rich repeat kinase in protection of dopaminergic neuron survival"

### Figure1_SourceData1

Figure 1-I
LRRK1 WB

control
LRRK1 KO

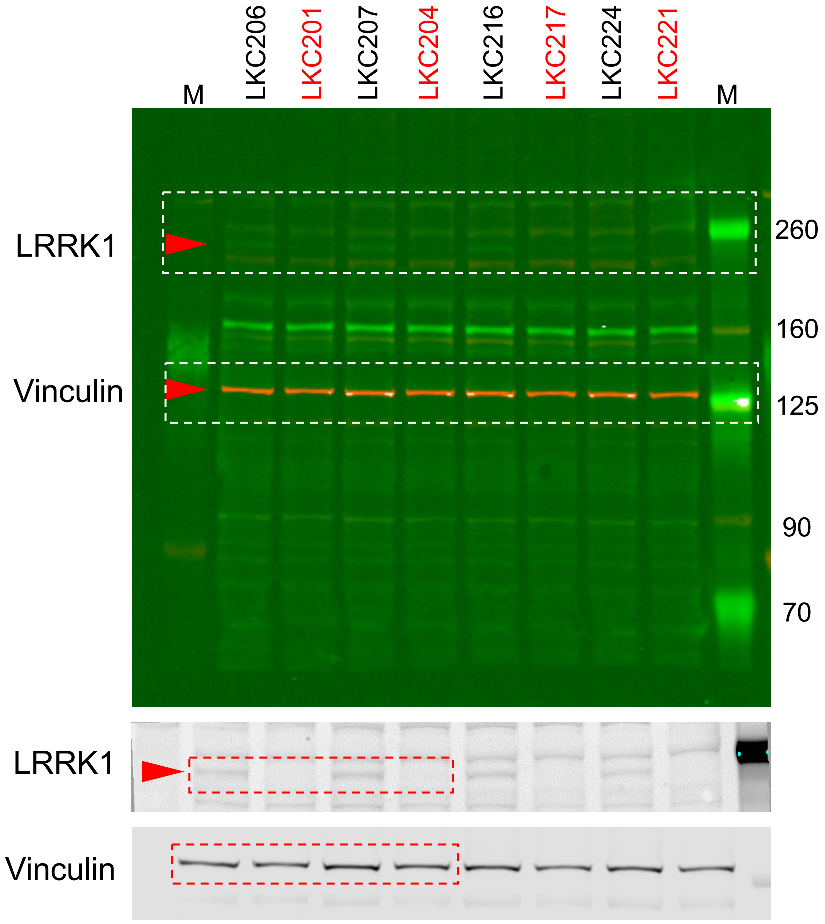


Figure 1-I
LRRK2 WB

control
LRRK2 KO

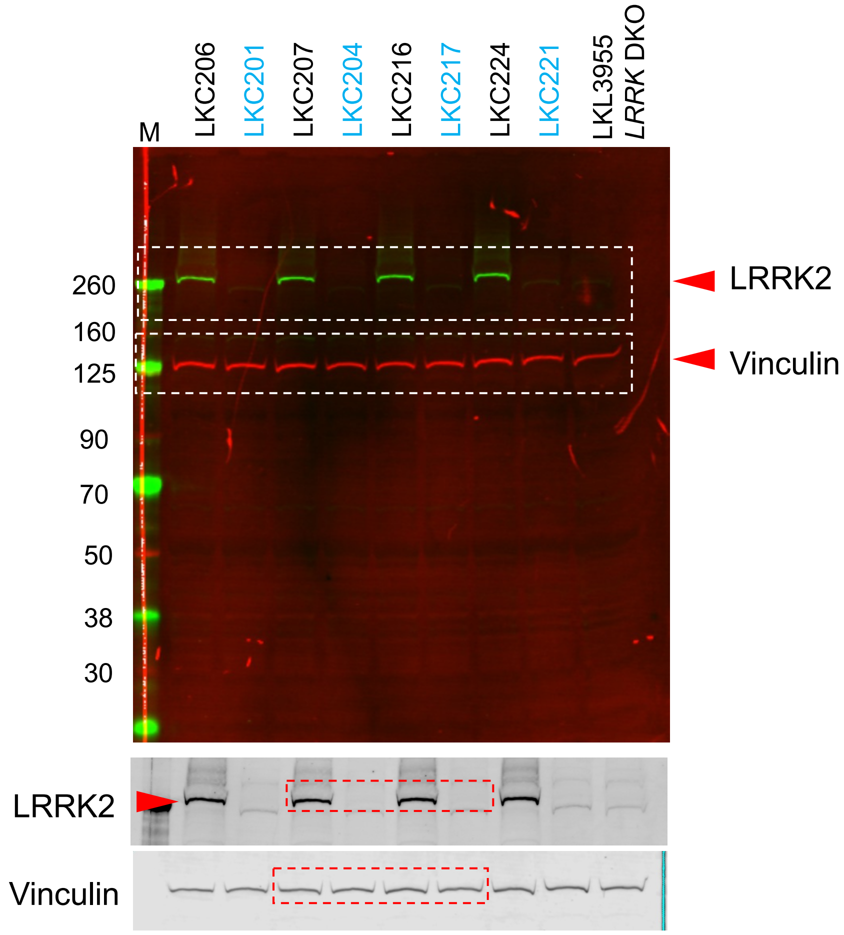
